# Golden eagles regularly use gravity waves to soar in the Alps: new insights from high-resolution weather data

**DOI:** 10.1101/2024.10.29.620992

**Authors:** Tom Carrard, Elham Nourani, Lukas Jansing, Tim Zimmermann, Petra Sumasgutner, Matthias Tschumi, David Jenny, Martin Wikelski, Kamran Safi, Michael Sprenger, Martina Scacco

## Abstract

Soaring flight developed as a result of behavioural and morphological adaptations that allow birds to reduce the metabolic cost of flight by harnessing the energy available in the atmosphere. Despite an increased attention given in the last decades to the physics and ecology that allow soaring flight, its study has been limited by the generally low spatio-temporal resolution of available atmospheric data. This constrained our ability to quantify the atmospheric conditions that allow soaring, and limited our understanding of its flexibility in different uplift conditions. While the use of updraughts such as thermals and orographic lifting are well described in the literature (albeit only quantified through atmospheric proxies), the use of others, such as gravity waves, was hypothesised but largely undocumented. Recent advancements in high-resolution atmospheric modelling, with hourly output available at the kilometer-scale grid spacing, offer new opportunities to investigate the flexibility of soaring flight in response to complex atmospheric dynamics. In this study, we used a combination of a high-resolution atmospheric analysis and high-resolution GPS tracking data to characterise the updraught sources used by golden eagles, *Aquila chrysaetos*, in the European Alps. We document that golden eagles in this region repeatedly use gravity waves, and that while thermals were still the main updraught source used for soaring, gravity waves were involved in at least 19% of the inspected soaring segments. In winter, when thermals were more scarce, the quasi-totality of soaring events were powered by gravity waves or orographic lifting, largely expanding the environmental energy available to soaring birds and therefore the landscape connectivity in topographically complex regions. Our results also emphasise the difficulty to distinguish between convective (thermals) and dynamic updraught sources, as these co-occur within the boundary layer over complex terrain.

## 2 Introduction

Flying animals across taxa have evolved different strategies to extract energy from the atmosphere and minimise their flying cost. In the last decade, the definition of the “energy landscape” as the environmental-driven variation in movement cost, and its role in shaping animal movement, have increasingly been recognised and integrated in movement ecology and landscape connectivity analyses, particularly in the aerial domain (Bohrer et al. 2012, Shepard et al. 2013, Katzner et al. 2015, Shamoun-Baranes et al. 2016, Santos et al. 2017, Péron et al. 2017, Williams & Safi 2021, Scacco et al. 2023). Early observational and theoretical studies already contributed to understanding the different atmospheric processes used by landbirds to fly at low energy costs (Pennycuick 1972, 1975, 1989, Kerlinger 1989, Shepard et al. 2016). Among these, thermals and orographic lifting are identified as the primary sources of atmospheric energy employed by large landbirds to soar. Thermals are small regions of ascending air generated by differential heating of the Earth’s surface, while orographic lifting results from the deflection of horizontal winds by hills and mountain ridges (Kerlinger 1989, Brandes & Ombalski 2004).

The movement decisions of soaring birds are known to be affected by the distribution of these uplifts, and many species adapted to extract energy from different uplift forms depending on their availability (Bohrer et al. 2012, Katzner et al. 2015, Péron et al. 2017). Because of the importance of uplift availability in determining the connectivity of landscapes at different scales, several studies have sought to quantify the contributions of various updraught sources to examine their differential use among species, age groups, and seasons(Katzner et al. 2015, Shamoun-Baranes et al. 2016, Duerr et al. 2015, Katzner et al. 2012, Murgatroyd et al. 2018). Many of these studies were driven by the need to understand birds energy budget in order to predict local and long-distance connectivity (e.g. migratory route selection) (Nourani & Yamaguchi 2017, Nourani et al. 2024, Brønnvik et al. 2024), but also potential conflicts between soaring landbirds and anthropogenic infrastructures, such as wind farms (Péron et al. 2017). Such modelling efforts and risk assessments require a better understanding of how atmospheric conditions shape the energy landscape and movement behaviour of soaring birds.

Typical atmospheric analysis products are unable to resolve the highly transient nature of turbulent boundary layer flows over complex terrain. This spatio-temporal resolution mismatch between atmospheric analysis data and bird tracking data (often available at sub-second resolution (Bohrer et al. 2012, Santos et al. 2017, Scacco et al. 2019)) have hindered our ability to investigate the complex atmospheric conditions encountered by flying birds. Consequently, many studies have relied on proxies derived from coarse-resolution atmospheric data (e.g. the widely used ERA5 reanalysis data, Hersbach et al. (2020)) to estimate updraught availability and analyse its effect on flight parameters at different scales. Although this approach strongly contributed to building our understanding of the energy landscape of large soaring birds, the developed models still resulted in substantial unexplained variance (Santos et al. 2017, Bohrer et al. 2012, Murgatroyd et al. 2018, Kerlinger 1989).

Recent advancements in high-resolution meteorological modelling have enabled atmospheric representations at scales of a few kilometres (e.g. Benjamin et al. 2016, Skamarock et al. 2008, Zängl et al. 2015). While this resolution remains insufficient to fully capture conditions at the spatio-temporal scale experienced by a bird in flight, it may provide sufficient detail to identify the types of updraughts that fuel soaring, and fill part of the spatio-temporal gap between the atmospheric information provided by these models and the behavioural information recorded by the bio-logging technology. In particular, some of these models have a resolution that is fine enough to explicitly resolve the physical mechanism of deep convection, which is otherwise parameterised in lower-resolution models. Their higher resolution also allows for a better representation of topography, leading to a more accurate modelling of local weather phenomena such as mountain waves (Baldauf et al. 2011). For example, Garstang et al. (2022) used a high-resolution model to illustrate that golden eagles (*Aquila chrysaetos* ) can exploit atmospheric gravity waves to sustain soaring flight during migration. Although gravity waves are well-known by sailplanes pilots (Kerlinger 1989), they have never been explicitly considered when modelling the energy landscape of soaring birds. Over complex terrain, such mountain waves occur when the stably stratified flow is deflected upward by an orographic obstacle, forcing it out of its equilibrium position (e.g. Durran 1990). The atmospheric response depends on the height and width of the mountain, as well as on the ambient atmospheric profiles. Under certain conditions, horizontally propagating lee waves can form (e.g. Whiteman 2000). Since the European Alps are largely east-west oriented, mountain gravity waves are particularly favoured when the large-scale flow has a strong meridional component, such as during south-foehn events (e.g. Jansing et al. 2022).

In this study, we used the unique combination of high-resolution weather analysis data and high-resolution Global Positioning System (GPS) data from a golden eagle in the Alps to provide a fine-scale quantification of the atmospheric conditions experienced by golden eagles flying in this region, while also investigating the capability of numerical weather models to distinguish between the different updraughts sources used by these birds. By integrating these two aspects we were able to document the eagles’ use of gravity waves compared to thermals and orographic lifting, and examined the use of the three updraught sources in relation to their temporal availability. To illustrate the steps that guided our investigation, we first present a case study of an eagle using a gravity wave and explain how a few visualisations allowed us to exclude other possible updraughts types. Using the same method, we then attribute 150 randomly selected soaring segments to different updraught types (thermal, orographic lifting, gravity wave and mixed categories), describing their relative contribution and their different properties. Our integration of high-resolution behavioural and meteorological data offers a new understanding of the atmospheric conditions shaping soaring flight, with implications for modelling energy landscape and landscape connectivity in topographically complex regions.

## 3 Data & Methods

### 3.1 Meteorological data

The present study used operational analysis data provided by the Swiss national weather service MeteoSwiss (https://www.meteoswiss.admin.ch). For the time period considered, MeteoSwiss employed the COnsortium for Small-Scale Modelling (COSMO) model for high-resolution data assimilation and numerical weather pre-diction on a regional domain covering the greater Alpine region. The COSMO model solves a non-hydrostatic formulation of the governing atmospheric equations on a computational grid (Steppeler et al. 2003, Baldauf et al. 2011). Atmospheric processes on the subgrid-scale are represented by parameterizations. Until October 2020, observational nudging was applied to combine the first guess from the model with the observations (Schraff 1997). Thereafter, the local ensemble transform Kalman filter technique was used to produce the analysis (Schraff et al. 2016). The data is provided at hourly steps and with a horizontal grid spacing of one kilometre and on 80 vertical levels. Such high-resolution analysis data can be used for research on mesoscale atmospheric processes such as wind systems in the Alpine region (e.g. Jansing et al. 2023).

At the kilometre-scale resolution, COSMO explicitly resolves deep convection and provides a representation of the vertical velocity field. However, soaring birds may rely on very short and/or very local updraughts that are not captured by the spatio-temporal resolution of COSMO. For this reason, we did not rely exclusively on the vertical wind field but used a larger set of surface and atmospheric variables to characterise updraught types. For our analysis, we used pressure, temperature, and the three vector components of wind. We also extracted the following surface variables: surface sensible heat flux, slope angle, slope aspect and elevation. With this set of variables, we considered three proxies for thermal and orographic lifting occurrence.

As proxies for thermal formation, we used the surface sensible heat flux and derived the atmospheric stability of the atmosphere in the lower 200 metres above the ground. The surface sensible heat flux represents the direct exchange of heat between the ground and the lowest layer of the atmosphere and is proportional to a temperature gradient. A sensible heat flux from the ground to the atmosphere is a prerequisite for thermal updraught formation and has been used for this purpose in the derivation of thermal proxies (Bohrer et al. 2012). The atmospheric stability was computed as the rate of change of potential temperature with altitude. Potential temperature is a common variable in atmospheric science and corresponds to the temperature *T* that an air parcel at pressure *P* would reach if brought adiabatically to a reference pressure level *P*_0_ of 1000 hPa. It is computed as:

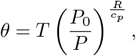

where *R* is the universal gas constant and *c*_*p*_ is the specific heat capacity at constant pressure. The vertical gradient of potential temperature gives a direct information on atmospheric stability and is positive for a stable stratification. A neutrally or unstably stratified atmosphere therefore indicates an environment where buoyant thermals are prone to form and rise due to density differences with respect to their surroundings.

As a proxy for orographic lifting, we computed a lifting coefficient proposed by Brandes & Ombalski (2004). The coefficient is then used to quantify the conversion from horizontal to vertical velocity by orographic lifting, as formulated by Bohrer et al. (2012):

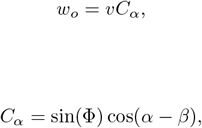

where *w*_*o*_ is a vertical speed, *v* is the horizontal wind speed, *C*_*α*_ is a lifting coefficient, Φ is the slope angle, *α* is the slope aspect and *β* is the horizontal wind direction. Negative values of *w*_*o*_ were set to zero as we are only interested in positive vertical velocities able to sustain soaring flight. Note that, because orographic lifting is represented in COSMO model and we are using the terrain characteristics as derived from the model orography, our proxy is merely an indication of how much the horizontal wind is aligned with the topography. A high value of *w*_*o*_ can indicate either a horizontal wind deflected by the topography or the occurrence of thermally induced upslope winds (Zardi & Whiteman 2013). This contrasts with other studies that have used *w*_*o*_ to approximate lifting of the horizontal wind by subgrid topography (Bohrer et al. 2012, Péron et al. 2017, Santos et al. 2017).

### 3.2 Golden eagle tracking data

We relied on existing tracking data from a long-term study of juvenile golden eagles in the resident population of Central European Alps and the North Apennines, spanning from 2017 to 2023. Eagle locations data were acquired via Global Positioning System by solar-powered transmitters manufactured by e-obs GmbH (Munich, Germany), fitted to the birds using leg-loop harnesses (for details on fieldwork procedure and sampling protocol see Zimmermann (2021)). The transmitters were set to record GPS data regularly every 20 min, but with fully charged battery, the loggers collected “super bursts” at intervals of 15 minutes. Super bursts consisted of 5 minutes of consecutive 1 Hz locations (1 location per second), and were the only data used in this study. The recording in bursts implies that data are irregularly spaced in time but with displacements being captured on a very fine scale. We selected the GPS track from 2020 and 2023 for which COSMO data were available.

To classify the updraughts source used by golden eagles, we selected a random set of 150 soaring segments from those available in 2023. We disregarded data from 2020 for this part of the analysis to avoid false conclusions due to potentially systematic differences in the weather data, as the assimilation method changed for 2023 compared to 2020. A soaring segment was defined as an uninterrupted sequence of points with positive vertical velocity. To filter out small-scale updraughts and turbulence, the vertical velocity values were smoothed with a 21 second moving window prior to segmentation. We excluded small ascents as we were interested in larger scale updraughts such as large thermals and gravity waves. We did this by setting a threshold of 234 metres height gain (difference between minimum and maximum height per segment). This allowed us to keep only the major climbing segments contributing to 50% of the total height gain of all segments.

### 3.3 Visualisation of weather conditions

In order to understand the atmospheric conditions experienced by the eagles in flight, and to characterise the updraught source used during soaring, we produced three types of visualisation. The first visualisation is a sequence of horizontal cross-sections, one for each point along the eagle trajectory. The cross-section plane was defined as a constant altitude plane at the elevation of the eagle GPS point. The weather variables were then interpolated to the plane and further linearly interpolated at the exact time of each GPS point (see later fig. 2).

**Figure 1.**
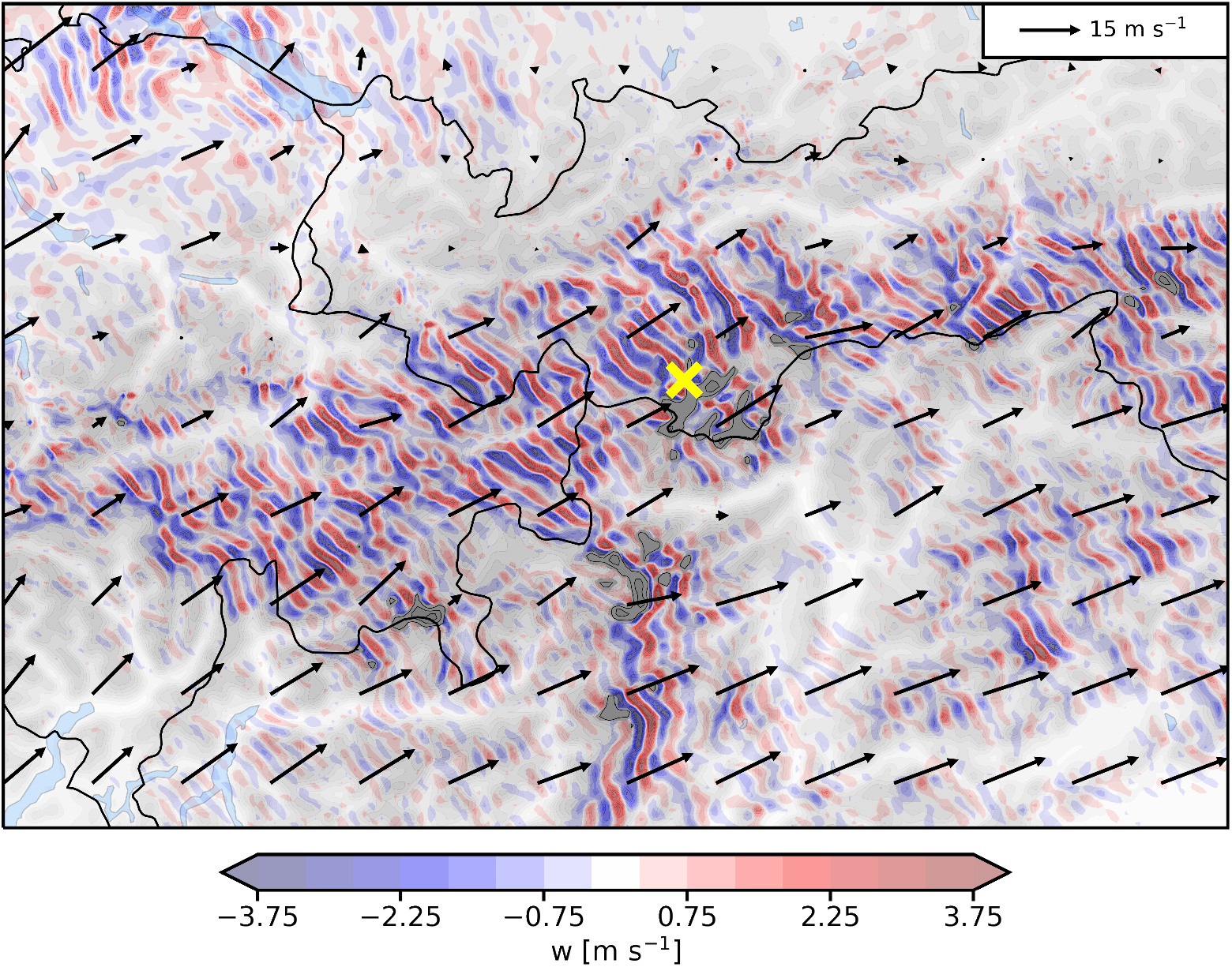
Geographical context of the case study on 21 October 2020. The location of the eagle track is indicated by the yellow cross. The background grey shading indicates the model topography. Red and blue transparent shading depict the oscillating vertical wind field (w) at 3000 m. Arrows show the horizontal wind vector at 3000 m altitude.

**Figure 2.**
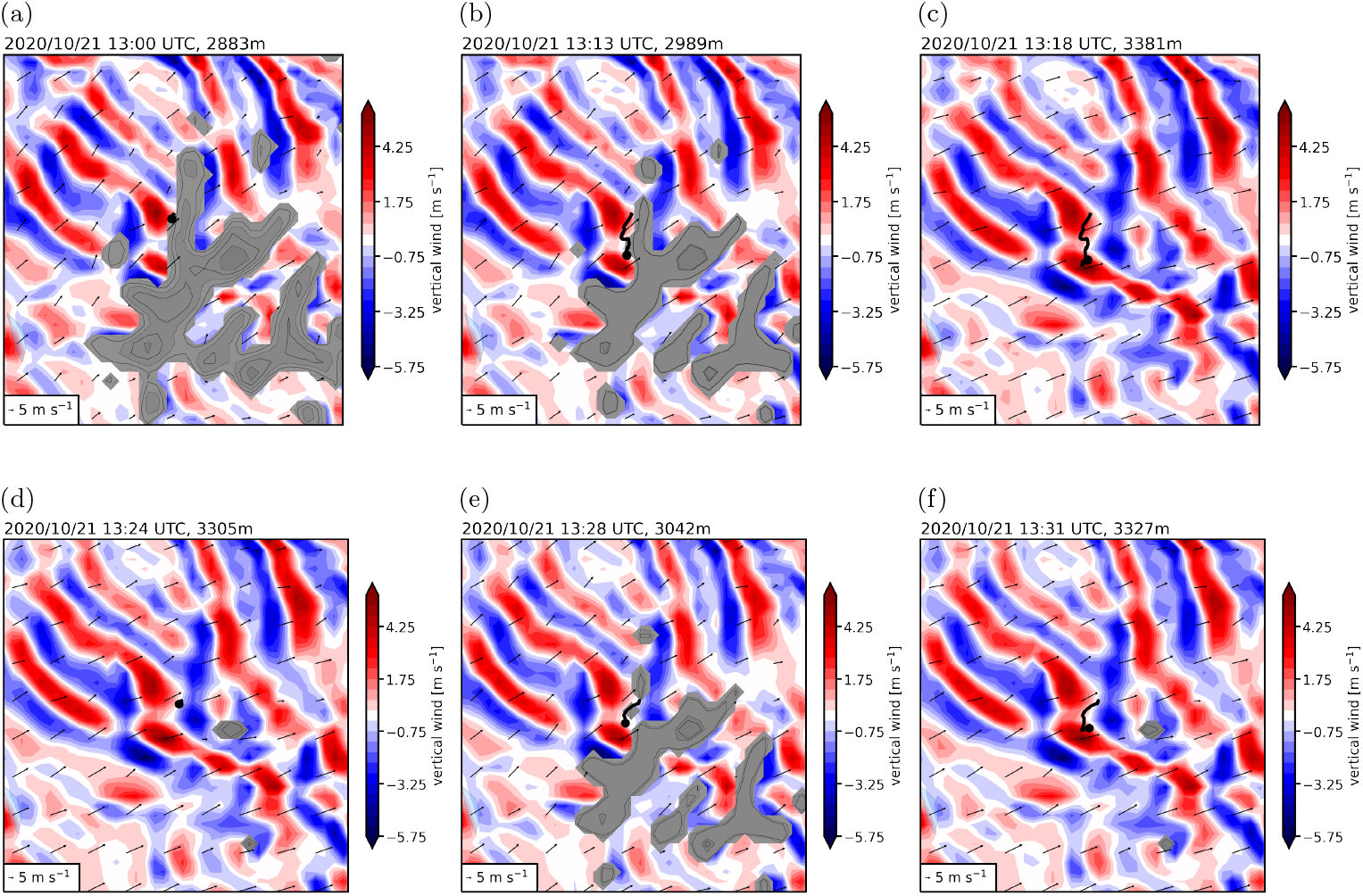
Horizontal cross-sections at the eagle elevation for two segments of gravity wave soaring. Coloured shading shows the vertical wind speed, grey shaded contours indicate where the cross section crosses topography and arrows indicate horizontal wind vectors. The black dots indicate the current location and the black line shows the track of the eagle from the start of the segment. The first row (a-c) corresponds to the three time steps of the first soaring event and the second row (d-f) to the second event, a few minutes later.

The second visualisation is a vertical cross-section. In contrast with the first visualisation, we considered a sequence of GPS points within an ascending segment. The vertical cross-section plane was then defined with the least square method to minimise the distance of the plane from all GPS points in the horizontal space (i.e. a linear regression on the longitude and latitude space of the GPS points). The weather conditions were then interpolated bilinearly to the vertical cross-section plane and further linearly interpolated in time at the average time of the sequence of GPS points (see later fig. 3).

**Figure 3.**
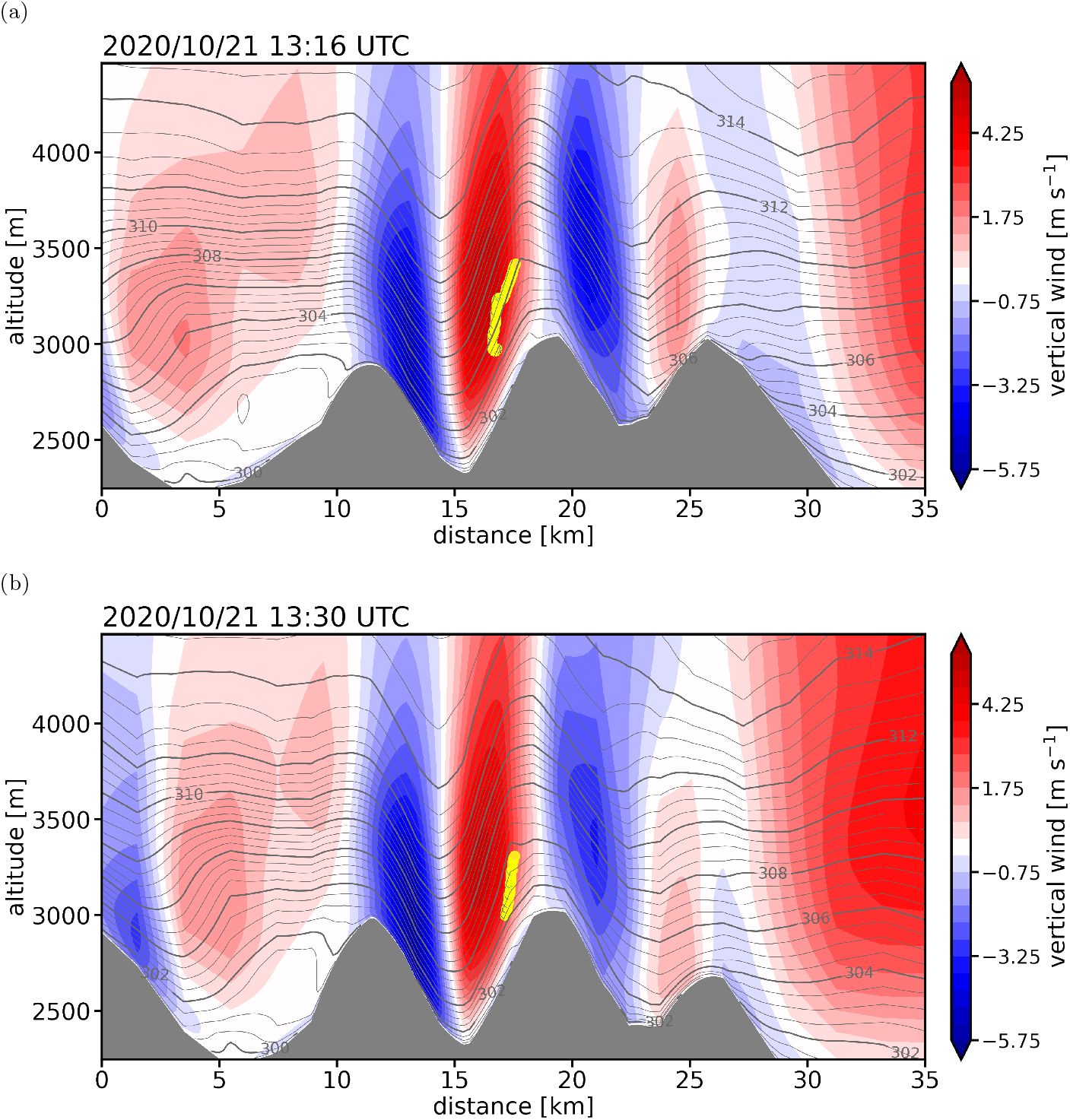
Vertical cross-sections corresponding to the two segments of gravity wave soaring in figure 2. The cross-section plane is defined to minimise horizontal distance to each location points of the eagle during the ascent (see section 3.3). Coloured shading shows the vertical wind field and grey shading indicates topography. Contours show lines of constant potential temperature. Yellow point indicate the sequence of locations of the eagle during the soaring segments.

We then computed a third visualisation to show the (interpolated) weather conditions at each eagle location along its track. For this, each model vertical level was interpolated to the eagle horizontal location and time. This yields a sequence of vertical profiles that can be arranged to yield a distance-height profile. The x axis represents the distance covered by the eagle from the start of the tracking segment and the y axis represents the altitude above sea level. In addition, this visualisation includes a lower panel showing the three chosen updraught proxies interpolated at the eagles location: surface sensible heat flux, lower layer stability and orographic uplift proxy (see later fig. 4).

**Figure 4.**
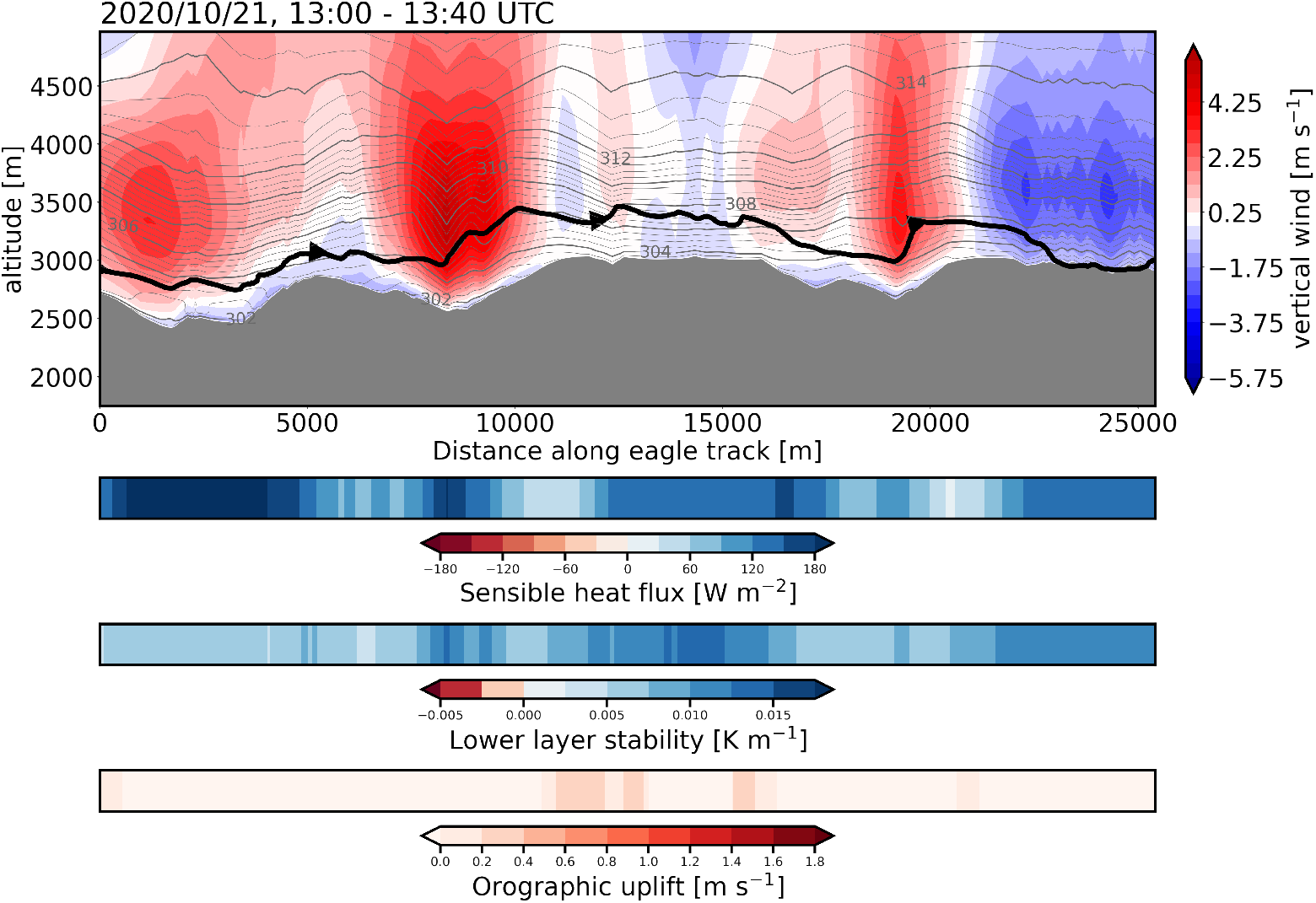
Full trajectory of the golden eagle with interpolated atmospheric conditions at each point of the trajectory. Coloured shading shows the vertical wind speed, grey shading indicates topography, and grey contours are lines of constant potential temperature. The thick black line indicates the trajectory of the eagle (larger arrows indicate flight direction). The lower panels show the three surface proxies interpolated at the eagle location.

### 3.4 Updraughts classification

To assess the relevance of including gravity waves as a potential updraught source, we first explored the entire data set (2020 and 2023) to find examples of gravity waves used for soaring. As gravity waves mostly occur when strong across-ridge winds are present (Wolff & Sharman 2008, Díaz-Fernández et al. 2021), we excluded all GPS bursts for which the wind speed at a pressure level of 700 hPa (around 3000 m) was weaker than 8 ms^−1^. For all bursts above this wind threshold, we then inspected the weather conditions at the locations of the eagles with the help of the visualisations described in section 3.3, and manually selected the cases where an eagle was flying in a gravity wave updraught. Waves were identified visually as alternating positive and negative vertical velocities on a vertical or horizontal cross-section (see later figures 1, 2, 3, and 4).

In a second step, we inspected the 150 randomly selected soaring segments from 2023 and attributed each segment to one of the following updraught categories: thermal, orographic lifting, or gravity wave. In terms of physics, leeward gravity waves are a consequence of windward orographic lifting. Because the gravity waves can propagate in the lee of a mountain ridge, the wave-induced uplift is not necessarily in phase with the topography. This means that updraughts due to gravity waves are not necessarily captured by the orographic lifting proxy widely used in the literature (Bohrer et al. 2012, Péron et al. 2017, Santos et al. 2017). In addition, since they form on the lee side of the mountain, gravity waves updraughts are potentially more challenging to reach and use than windward orographic lifting updraughts (Kerlinger 1989). For these reasons, we considered gravity waves and orographic lifting as two distinct updraught categories in the following classification.

The segments were classified based on the visualisations described in the section 3.3. We used the weather conditions and the surface proxies to classify each segment based on our understanding of atmospheric updraught. Sometimes, the conditions were indicative of two different types of updraught. In these conditions we labelled the segments as a mixed category (e.g. thermal/orographic lifting or thermal/wave). Note that following our reasoning in the last paragraph, we did not allow for mixed category of orographic lifting and gravity wave events, despite they can occur in combination in the atmosphere. In some other cases, we were not able to determine the updraught source based on the weather conditions from COSMO and we classified the segment as ‘unknown’. An example of the criteria used to classify the segments can be found in sections 4.1 and 4.2 and Appendix **??**.

### 3.5 Updraughts characteristics

To assess the consistency of our labelling approach, we performed two principal component analyses (PCA) on our sample of soaring segments. For each of the 150 segments, we computed the mean surface sensible heat flux, mean stability in the lowest 200 m, mean orographic lifting proxy, mean horizontal wind speed, and, as a rough proxy for boundary layer depth, the mean height of maximum stability. To assess the impact of the different updraughts on flight characteristics, we also calculated for each segment the total altitude gain, mean vertical speed, maximum and mean height above ground, total horizontal distance and total absolute turning angle (computed as the sum of the absolute difference in heading between two consecutive segments). For each set of variables, weather variables and flight variables, we then performed the principal component decomposition to explain at least 90% of the total variance.

## 4 Results

### 4.1 Case study: Gravity wave soaring

We first present a case study on a gravity wave soaring event occurring on 21 October 2020. On this day, the synoptic-scale weather was dominated by an upper-level trough over the eastern Atlantic, inducing a strong southwesterly flow in the Alpine region. This flow configuration corresponds to a deep-foehn situation over the Alps and is prone to gravity wave activity (Jansing et al. 2022). The eagle track was recorded in the Austrian region of Tyrol in the vicinity of the Swiss and Italian borders (fig. 1) at an elevation of about 2800 m.

Horizontal cross-sections (fig. 2) show a clear wave pattern in the vertical wind field. At 13:13 UTC (fig. 2b), the eagle first reaches a region of strong vertical updraught and gains about 500 metres in six minutes with a maximum vertical speed of 7.6 ms^−1^. The bird then glides back to its original position (not shown) before returning to the same gravity wave updraught at 13:28 UTC (fig. 2e) where it climbs about 350 metres in four minutes with a maximum vertical speed of 6.4 ms^−1^. Each of these two ascents is pictured in a vertical cross section in figure 3. The modelled gravity wave updraught has a depth of more than 2000 metres with a stable stratification (regularly spaced contour lines of potential temperature with a positive vertical gradient, fig. 3), which is a prerequisite for gravity wave formation (Whiteman 2000).

Figure 4 shows the full trajectory of the eagle with the local conditions encountered by the eagle at each point of its trajectory. The updraught proxies shown in the lower panel indicate that thermal updraughts cannot form because the atmosphere is strongly stratified and the surface sensible heat flux is positive. Note that the surface fluxes in COSMO are positive for downward fluxes by convention. In addition, orographic lifting is not likely, as the orographic uplift proxy shows that the horizontal wind at the surface is not aligned with the slope (low values). This indicates that the updraught is probably caused by the horizontal propagation of the gravity wave on the lee side of the relief rather than by local lifting of the horizontal flow on the windward side. However, as the gravity wave is phase with the topography, local lifting cannot be totally excluded in this case study, emphasizing that gravity wave and orographic lifting cannot always be safely distinguished. The two large updraught regions (red shading in the figure, right before 10 km and 20 km) correspond to the same gravity wave updraught visited twice by the eagle, as seen in figure 2.

This latter point is very important as it could be argued that the eagle is flying accidentally in the gravity wave. The combination of figures 2 and 4 clearly shows that this is not the case, and that the eagle is revisiting the stationary updraught to exploit the strong vertical wind, suggesting that mountain waves are another type of updraught that can be exploited by golden eagles. While this case study clearly shows that gravity waves, in agreement with the work from Garstang et al. (2022), can be used for soaring flight, their contribution and ecological importance for golden eagles flight are yet to be explored. In the next section, we bring additional quantitative evidence that gravity wave updraughts should be accounted for when studying bird soaring flights.

### 4.2 Frequency of updraughts use

Figure 5 shows the location of the 150 random segments selected to investigate the different types of uplift used by golden eagles. Most segments are located in the Italian Alps, with a few in Austria and Switzerland, five in the northern Apennines in Italy and one single event in a flat region close to the coast. Of the 150 segments, 19 occurred from December to February, 62 from March to May, 52 from June to August, and 17 from September to November. The irregular monthly distribution of the soaring segments reflects the sampling frequency of the GPS, which depends on the battery level and therefore on sunlight availability.

**Figure 5.**
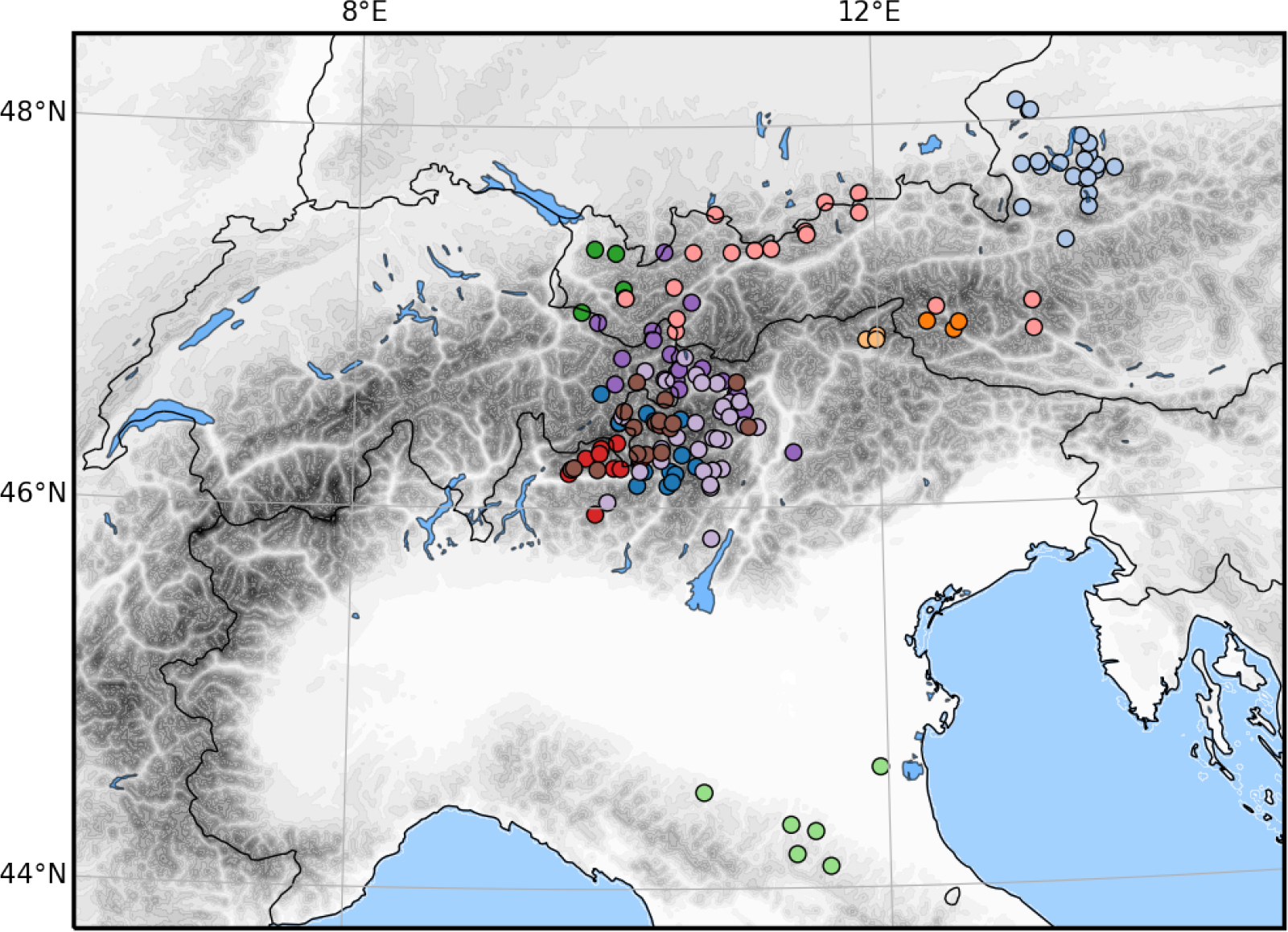
Average location of the 150 randomly selected soaring segments. Each colour corresponds to a golden eagle individual (11 individuals for the 150 segments). The model orography is shown in the background for orientation.

Three examples of visualisations used to label the updraughts are presented in figures 6, 7, and 8. In figure 6, as the vertical wind speed interpolated from COSMO is zero over the span of the soaring segment, the updraught must have been caused by small-scale processes not explicitly resolved in the analysis data. In addition, the horizontal wind is relatively weak and the orographic lifting proxy is close to zero, making dynamically induced updraughts unlikely. Instead, conditions prone to the formation of thermals can be identified by the high (negative) surface sensible heat flux values and the low stability in the lowest 200 metres of the atmosphere. Consequently, this example of a soaring segment is classified as thermal updraught. In the contrasting second example, the orographic lifting segment is characterised by a high orographic uplift proxy and high horizontal wind speeds at higher levels (fig. 7). In this case we can exclude a thermal contribution due to the sensible heat flux values close to zero. Furthermore, the vertical wind field is slightly positive in a narrow band close to the surface, which suggests that COSMO partly captures the vertical deflection of the wind by the topography resulting in a slope flow.

**Figure 6.**
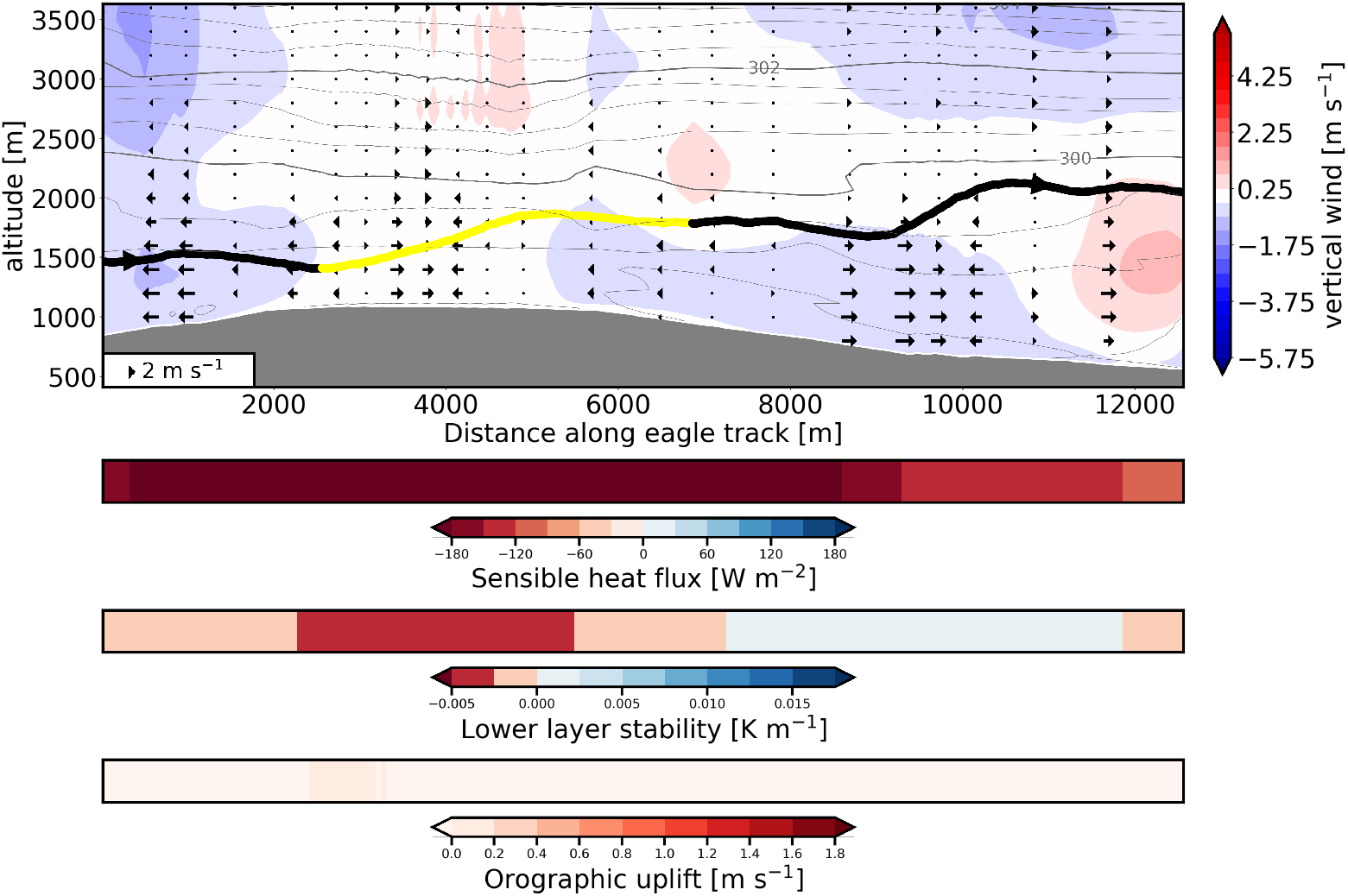
Example of a soaring segment classified as thermal updraught. The figure depicts the vertical profiles of vertical wind speed (shading), potential temperature (grey contours) and the horizontal wind projected along the eagle trajectory (arrows). The grey shading represents topography. The lower panels show the sensible heat flux (negative values for an upward flux), lower layer stability (potential temperature gradient in the lower 200 m), and orographic lifting proxy. The black line shows the eagle track and the yellow part corresponds to the labelled soaring segment (larger arrows show the direction of flight).

**Figure 7.**
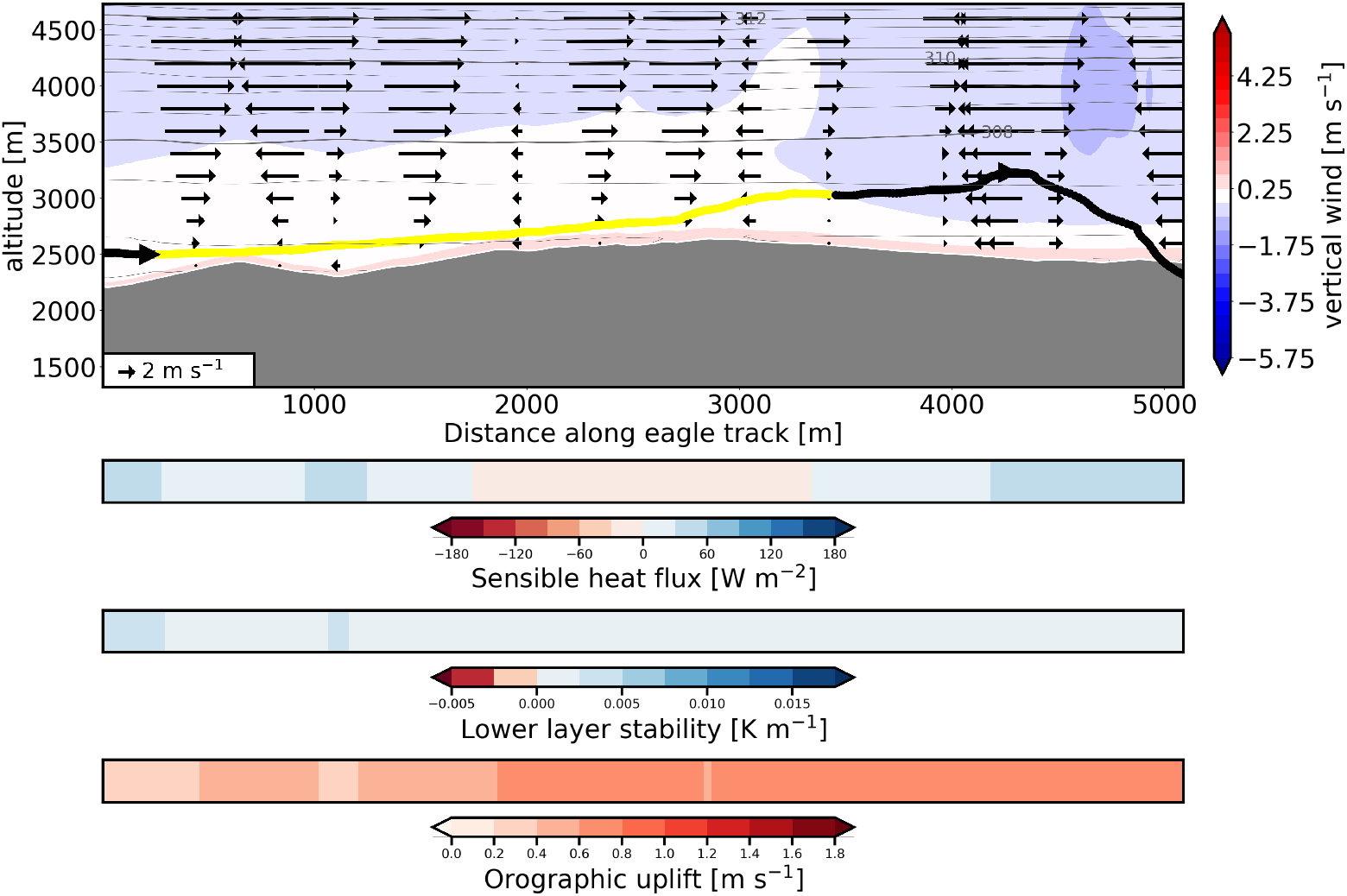
As in figure 6, but for an updraught induced by orographic lifting. Note that the eagle track intersecting the topography at the end of the track is either a result of a GPS error and/or an inconsistency of the interpolation (the interpolated topography does not perfectly represent the true topography).

**Figure 8.**
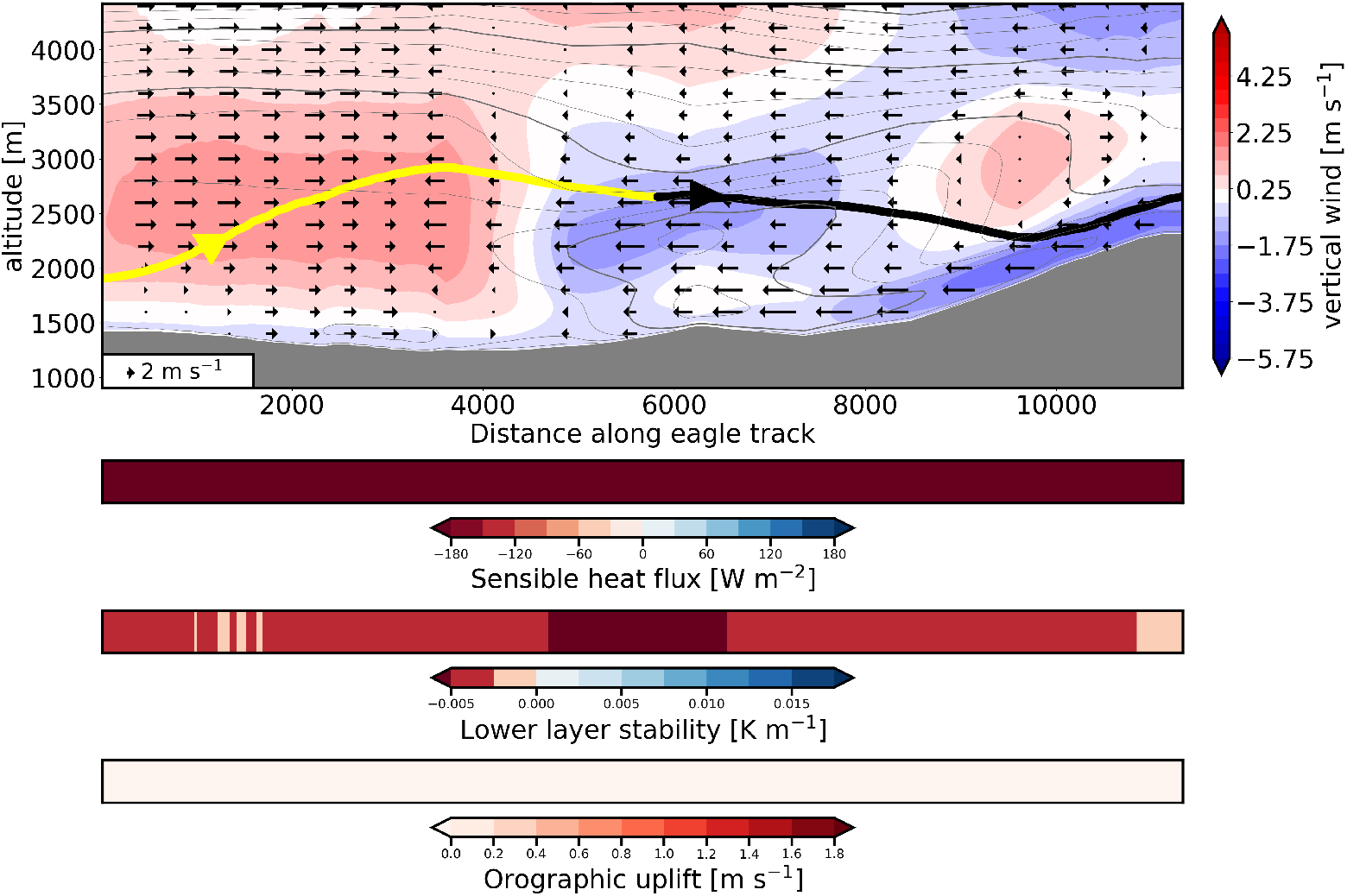
As in figure 6, but for combined thermal and gravity wave updraught.

Finally, figure 8 illustrates an example of combined updraught sources (mixed category), where the eagle’s flight is likely supported by both thermal and dynamic processes. The large region of positive vertical velocity detached from the ground corresponds to a gravity wave updraught, as can be confirmed by looking at an horizontal cross-section (not shown). The high stability above 3500 m altitude and the strong horizontal winds are consistent with gravity waves formation. In the first part of the segment however (before 2000 m distance), the eagle is flying in a region of low stability (the lines of potential temperature are far apart) and we observe high surface sensible heat fluxes towards the atmosphere throughout the segment. These conditions are prone to thermal formation and hint towards the occurrence of thermals in the lower atmosphere. Based on this visualisation, we conclude that the eagle is benefiting from both the low-level thermal updraught and the higher level gravity wave updraught. This example nicely illustrates how thermal and dynamic effects can co-occur in the atmosphere and questions the relevance of binary classifications.

With our workflow (see section 3), we were able to classify 77% of the 150 randomly selected segments. The remaining 23% of the segments were categorised as unknown (fig. 9b). Among the 115 successfully classified segments, 32% were classified as mixed events (i.e. thermal/gravity wave, or thermal/orographic lifting). Thermal updraughts were found to be the main source of energy for soaring flight in our dataset, contributing to nearly 60% of the selected segments (70% when disregarding the ‘unknown’ events) (fig. 9b). Orographic lifting contributed to 24% of the segments (32% when disregarding ‘unknown’ events). Gravity waves contributed in 19% of the selected events (24% when disregarding the ‘unknown’ events), suggesting that they represent an important source of energy that is regularly used by golden eagles. By summing up the contribution of both orographic lifting and gravity waves, we obtain a total contribution reaching 56% of all cases. This means that dynamically induced updraughts contributed to more than half of the soaring segments considered in this study.

**Figure 9.**
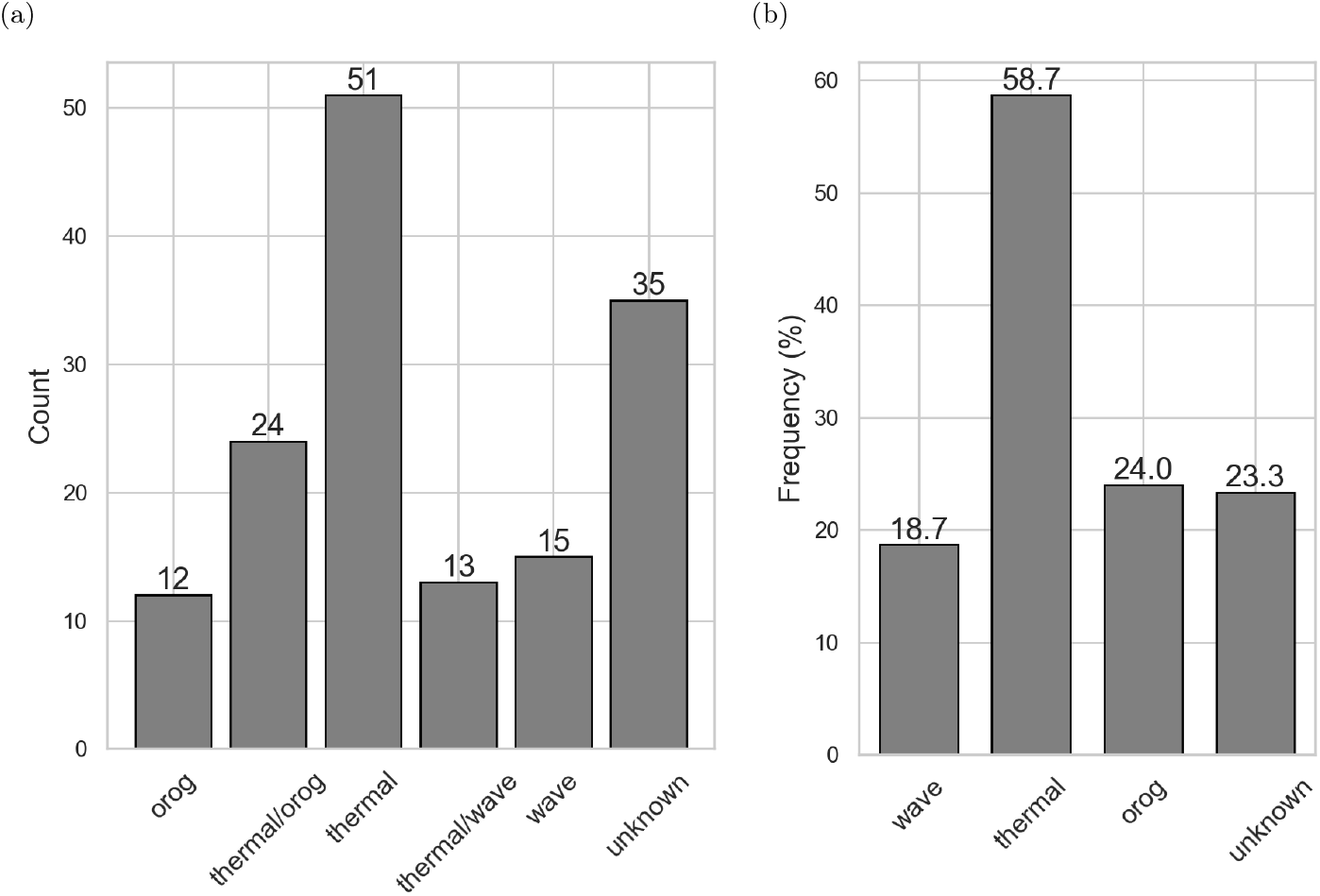
Result of the classification of the 150 randomly selected soaring segments. a) shows the the raw number of segments for each updraught category while b) shows the relative contribution of each updraught type. In b), each column contains the number of segments of the respective updraught type + the corresponding mixed events. This means that the ‘wave’ bar contains both ‘wave’ and ‘thermal/wave’ events from a) (i.e. 28 events divided by 150 total segments), such that the sum of all percentages exceeds 100%. Note that ‘orog’ and ‘wave’ stand for orographic lifting and gravity wave, respectively.

The monthly distribution of updraughts sources across the year exhibits a very clear seasonal effect, with the quasi totality of updraughts being dynamically induced (gravity wave or orographic lifting) from November to February, and a majority of thermal updraughts from April to July (fig. 10). This is explained by the lower availability of thermal energy in winter due to weak insolation and increased snow cover over the Alps, which force the eagles to rely on dynamic updraughts. During summer months, dynamic and thermal effects often occur in combination. Despite a high variability in sample size across months, this seasonal trend also indicates that the overall frequencies displayed in figure 9a may vary greatly between seasons. Because our sample is not balanced between months, this also suggests that our overall result is likely to underestimate the importance of gravity waves and orographic lifting, which are the dominant energy sources during winter months.

**Figure 10.**
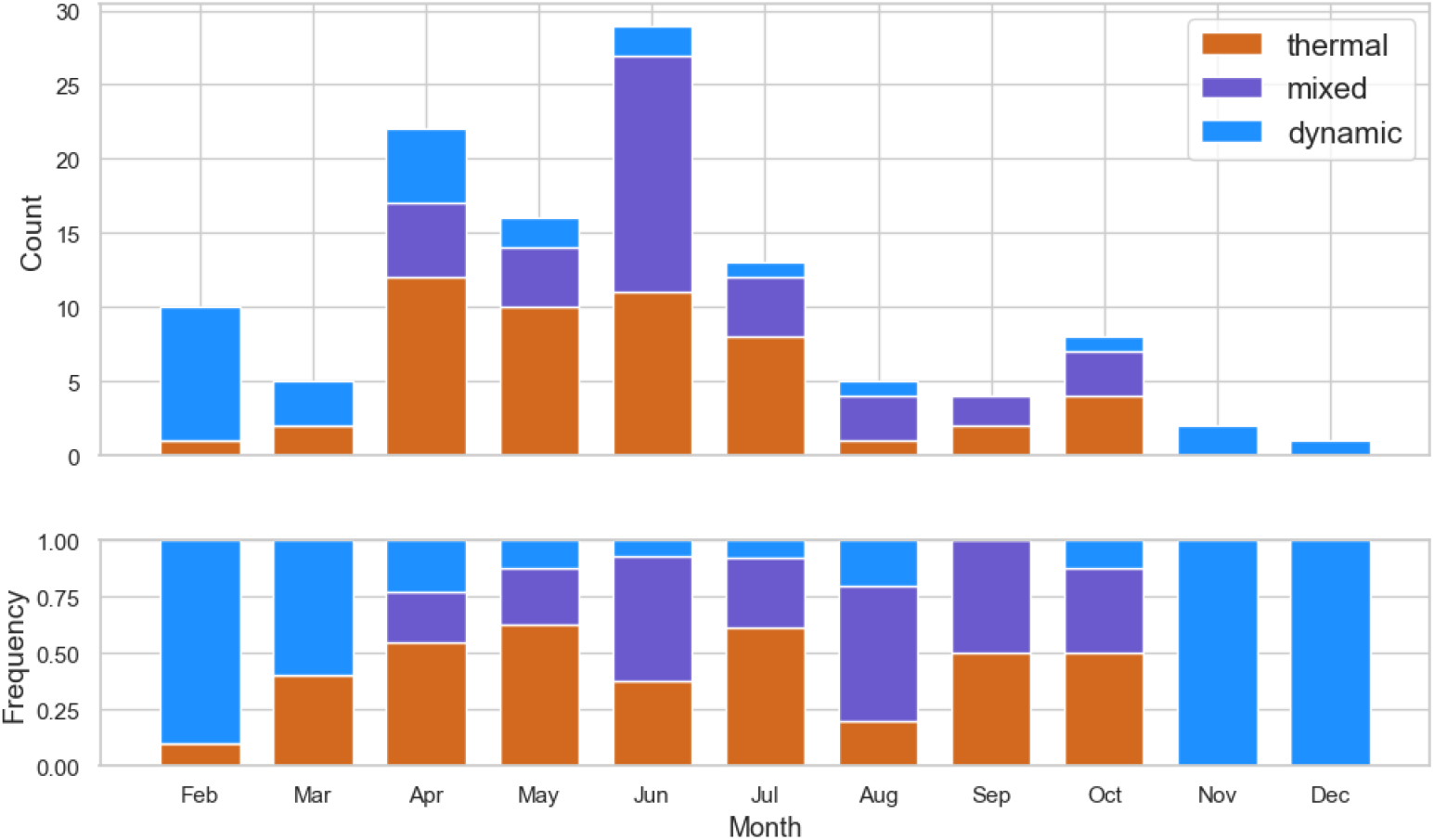
Monthly distribution of updraught types. Dynamic events (blue) include orographic and gravity wave updraughts and mixed events (purple) contain thermal/orographic and thermal/wave events. The upper panel shows the absolute number of segments per category while the lower panel displays the proportion of each category.

### 4.3 Principal component analysis

To assess the consistency and reliability of our updraught classification, in terms of both meteorological and flight characteristics, we performed two PCAs. The PCA based on meteorological variables explains 95% of the total variance based on its four first principal components (PCs). PC 1 (explaining 38% of variance) was mainly a combination of surface sensible heat flux and surface stability and helps distinguishing gravity waves from thermals (fig. 11a and 11b). PC 2 (explaining 25% of variance) was mainly a combination of wind speed and height of maximum stability and distinguishes between thermals and dynamic updraughts (fig. 11a and 11c). PC 3 (explaining 17% of variance) was mainly a combination of orographic lifting proxy and height of maximum stability and best separated orographic lifting from thermals and gravity waves (fig. 11b and 11c). Finally, PC 4 (explaining 14% of variance) was a combination of all variables and did not allow us to distinguish thermals, orographic lifting, and gravity waves. However, PC 4 represented the best dimension to separate mixed events from each other (fig. 11d).

**Figure 11.**
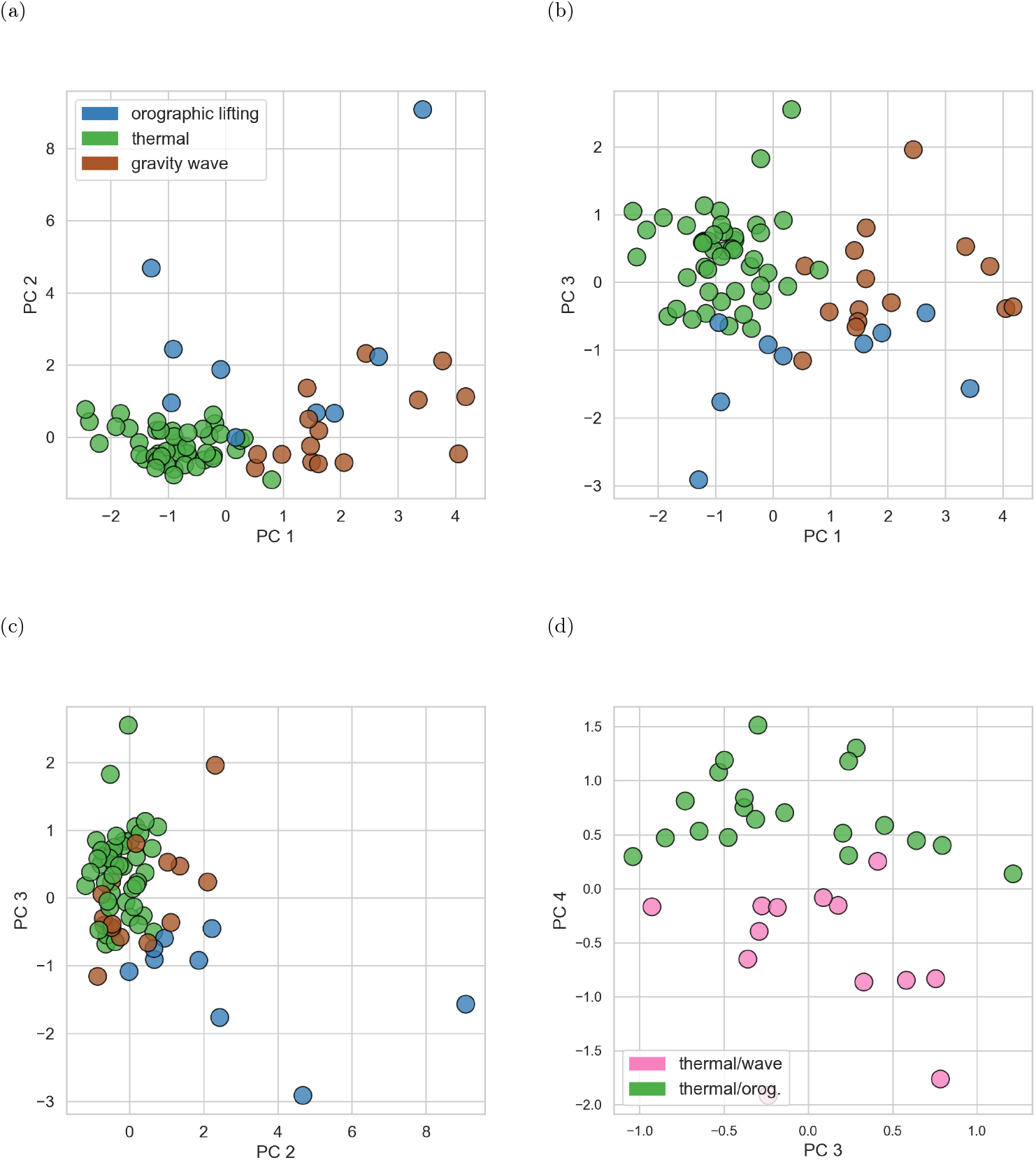
Labelled soaring segments plotted along the axis of the principal components analysis based on meteorological variables. The meteorological variables used for the PCA are surface sensible heat flux, stability of the lower 200 m, height of maximum stability, wind speed, and orographic lifting proxy.

The clustering of the different updraught types along the principal components extracted from the meteorological variables suggests that our classification is consistent in terms of physical variables. Although our classification demonstrates that it is difficult to classify updraughts into discrete categories (due to the possible combination of thermal and dynamic effects), this principal component decomposition indicates that we can define specific regions of the multi-dimensional phase space for which one updraught type is more likely than another. For instance, PC1 can be used to define a region where thermals are the most likely updraught type, or a region where thermals cannot occur. However, there is a region in between where both thermals and waves can co-occur and potentially be used in combination. This means that updraught types can continuously occur for a given range in the phase space of atmospheric variables and not only in discrete regions.

In the PCA based on flight behavioural variables (fig. 12), the three first PCs allowed us to explain 96% of the total variance. Flight behaviour alone, however, does not seem to be able to separate well the different updraught types 12, as the distribution of the selected variables does not seem to differ significantly between uplift sources. While we cannot exclude some level of correlation, this result shows no direct correspondence between flight behaviour and updraught type, suggesting that golden eagles can adopt a variety of flight strategies for a same updraught category. This also indicates that a weather-based classification of updraughts may yield very different results than a flight-based classification.

**Figure 12.**
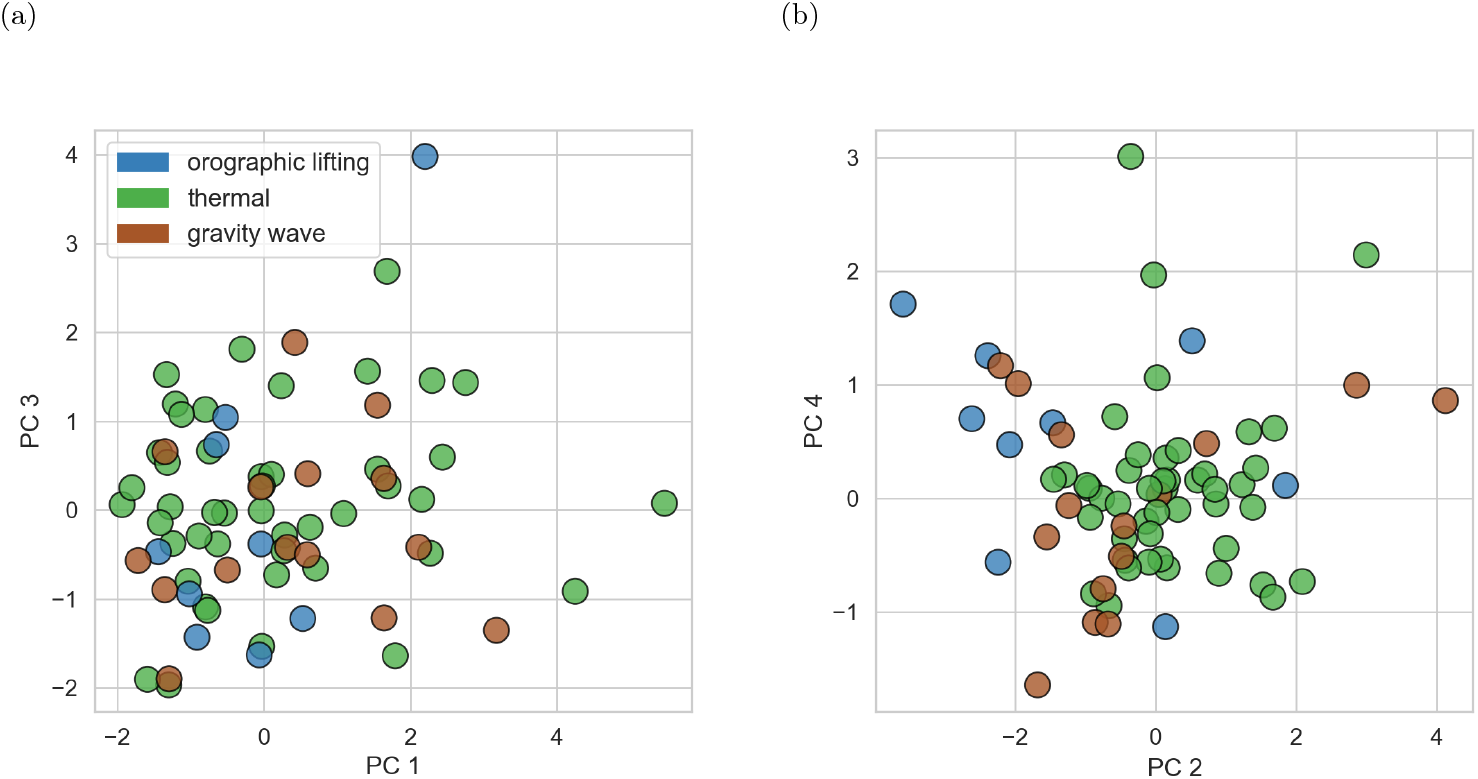
Labelled soaring segments plotted along the axis of the principal components analysis based solely on flight characteristics. The flight variables used for the PCA are total altitude gain, mean vertical speed, maximum and mean height above ground, total horizontal distance and total absolute turning angle.

## 5 Discussion and Conclusions

Through the combination of high-resolution movement data and weather information, this study offers a new perspective on the availability of environmental energy in topographically complex regions. By visualising high-resolution analysis data interpolated at the eagles’ GPS locations, we first demonstrated that gravity waves are being used by golden eagles as an energy source for soaring flight. Based on this observation, and in order to assess the importance of gravity waves for the soaring flight of golden eagles, we systematically inspected the weather conditions for a random set of 150 soaring events. In the majority of the cases (77% of the soaring segments), our approach allowed us to identify the updraught source used by the eagles and showed that gravity waves were involved in 19% to 24% of the segments.

This result constitutes the first robust evidence that golden eagles can use gravity waves to sustain soaring flight. While Kerlinger (1989) argued that gravity waves are the most powerful updraught that can be used to soar safely, he suggested that they were probably only rarely used by soaring raptors, because their properties make them both difficult to reach and to soar within. Our analysis not only indicates that gravity waves are a significant updraught source for golden eagles, but that together with orographic lifting, they could be the main source of energy supporting soaring flight during the winter months (fig. 10) and possibly during the hours on a day where thermals are less abundant. In addition, as the GPS tracking devices are solar-powered, our quantification of dynamic updraught contribution is possibly underestimated, and biased towards thermal updraughts. This is because the GPS tracking devices are more likely to record high-resolution data when solar energy is available, and solar energy is directly impacting thermal updraught formation. This means that the results displayed in figure 9a and 9b possibly underestimate the importance of dynamic updraughts which may occur more often when the GPS device is not recording. This is also consistent with the fact that dynamic updraughts were mostly detected in combination with thermal updraughts (fig. 9a), because they are less likely to be detected in isolation, when the conditions are not favourable for thermal formation.

In 32% of the cases, atmospheric conditions indicated that two different updraught sources were possibly contributing to the same soaring event (e.g. thermal effects were often observed in combination with orographic lifting). We think that this result reflects the complexity of the atmospheric boundary layer and that a simple binary classification between dynamic and thermal updraughts is often not possible nor realistic. Our current understanding of bird soaring in different updraughts has been largely influenced by our ability to observe it and to analyse it in combination with the available atmospheric information. While thermals are mainly described as broad columns of rising air occurring on sunny days with light winds, and orographic updraughts are considered to be winds vertically deflected by mountain slopes, our fine-scale analysis showed that it is often difficult to attribute real cases strictly to either of these two well-defined categories. These 32% of events classified as mixed events could represent part of the reason behind the high unexplained variance in soaring models developed in previous studies (Santos et al. 2017, Bohrer et al. 2012, Murgatroyd et al. 2018, Kerlinger 1989). In fact, by reducing the availability of updraught sources to a binary classification problem, these models may lack a representation of the inherent complexity of atmospheric processes.

Even though our small dataset did not allow for a systematic hypothesis testing, we observed that the updraught types classified by our approach could be clustered along the PC dimensions derived from the meteorological variables, but not along those derived from flight behaviour characteristics alone. This suggests that although some correlation between flight characteristics and updraught sources cannot be ruled out, their relationship is more complex than previously thought. In fact, the widespread assumption in the literature that slope or linear soaring - often described as gaining height in a straight line without circling - is fuelled by orographic lifting (Katzner et al. 2012, 2015, Williams et al. 2015, Santos et al. 2017, Murgatroyd et al. 2018), ignores thermal effects and anabatic winds that could contribute to the same soaring event. In the same way, dynamic lifting (orographic uplift and gravity waves) can create high-reaching updraughts that can allow birds to fly vertically without necessarily following the slope. This was nicely exemplified with our gravity wave case study, where the deflection of the horizontal wind led to a deep updraught region and a steep soaring ascent of the eagle. This methodological difference (i.e. between flight-based and weather-based classifications) may partly explain why we detected a larger proportion of dynamic updraughts compared to former studies (e.g. Katzner et al. 2015). We thus think that using a behavioural-based classification of updraughts types underestimates the importance of dynamic updraughts such as windward slope lifting and gravity waves. This finding, together with the seasonal distribution of updraughts sources shown by our results, also offers a new perspective on the flexibility of soaring flight. Apparently, soaring animals can adapt to an extremely dynamic environment and are able to extract energy from the different atmospheric processes they encounter to reduce their flight cost. Nonetheless, as we only used a small set of flight variables and a relatively small sample, a more in-depth analysis of the link between flight characteristics and updraught type would be required to confirm that they are unrelated.

In addition to revealing the use of gravity waves by golden eagles, our study highlights the benefit of combining high-resolution weather variables with high-resolution behavioural data to provide a three-dimensional representation of the aerial environment experienced by soaring birds that is much more detailed than what has been previously possible. The documentation of gravity waves and the assessment of their importance relative to other updraught sources was made possible by the use of high-resolution weather analysis data resolving mountain waves at a spatio-temporal scale that is relevant for golden eagles soaring flight. A finer resolution is particularly helpful over complex terrain such as the Alps, as the small-scale variability of orography and surface characteristics directly impacts thermal and dynamic effects, which determine the availability of atmospheric updraughts. This implies that despite atmospheric updraughts are still not fully resolved at the scale at which a flying bird experiences them, the use of surface characteristics and advanced visualisation can help identify the atmospheric processes involved in sustaining soaring flight. In this regard, the increased availability of high-resolution numerical weather models is promising for our understanding and modelling of the energy landscape of flying animals. Indeed, our novel approach has also allowed the creation of a classified dataset that can serve as a benchmark for predicting updraught types in larger golden eagle GPS datasets. As high-resolution atmospheric data are surely more representative of the actual conditions experienced than proxies derived from coarser models, this approach could significantly improve our representation of the energy landscape.

Understanding the energy landscape of soaring birds has been the focus of many behavioural studies in the last decade, because of its implications for human-wildlife interaction, especially in the context of anthropogenic infrastructure such as wind farms, but also for habitat suitability studies focusing on predicting landscape connectivity. For all of the above, an accurate comprehension of the weather conditions that allow soaring flight is fundamental. The methodological approach we proposed, together with the interdisciplinary collaboration between meteorologists and behavioural ecologists fostered by this study, allowed us to move solid steps in this direction. In addition to systematically quantifying the use of gravity waves - a largely ignored energy source for soaring flight - our approach highlights that the complexity of the atmospheric processes relevant for soaring has probably been underestimated. We believe that our successful classification of the updraught sources used in 115 soaring events offers a new perspective on the energy landscape in topographically complex regions, with profound implications for the modelling of soaring flight and landscape connectivity.

## Authors’ contribution

TC MSC EN and KS conceived and designed the study, KS PS MT DJ and MW provided the tracking data, TC carried out the analyses, TZ LJ and MSP provided analytical support, TC MSP and MSC interpreted the results and wrote the first draft of the manuscript. All authors contributed suggestions and text to subsequent drafts. All authors gave final approval for publication.

## Acknowledgements

We thank MeteoSwiss for providing the COSMO-1 analysis data. We are grateful to Martin Grüebler, Julia Hatzl and Enrico Bassi for their golden eagles’ data contribution. Andrea Roverselli and Klaus Bliem for their support during the eagles’ tagging. We are grateful for the crucial assistance of the mountain rescue teams of the Guardia di Finanza as well as the forestry and game wardens of South Tyrol/Alto Adige and the national park team in Gesäuse, as well as the Stelvio national park Italy and the Fish and Game Department of the Canton of Grisons (AJF) and many gamekeepers. We also thank the accompanying veterinarians Drs. Michel Mottini and Gilberto Volcan for their time and expertise in the field.

## Ethics

The handling and ringing of golden eagle nestlings in Switzerland was carried out under the authorization of the Office for Food Safety and Animal Health (ALT) of the canton Grisons (licence No. GR2017 06, GR 2018 05E, GR 2019 03E). In Italy, the permissions for handling, tagging and marking were obtained from autonomous region of South Tyrol (Dekret 12257/2018 and Dekret 8788/2020), as well as from the region of Lombardia for ringing and tagging through in Lombardia and South Tyrol by ISPRA (Istituto Superiore per la Protezione e la Ricerca Ambientale) with the Richiesta di autorizzazione alla cattura di fauna selvatica per scopi scientifici (l.r. 26/93). In Austria, all procedures for handling, tagging and marking were approved by the Ethics Committee of the University of Vienna (No. 2020-008) and permitted by the Federal Ministry for Education, Science and Research (No. 2020-0.547.571), Styria (BHLI-165942/2021-2) and Upper Austria (LFW-2021-263262/7-Sr). Finally, in Germany birds handled, tagged and ringed were done so under the permission issued by the government of Oberbayern (2532.Vet 02-16-88 and 2532.Vet 02-20-86). All procedures followed the ASAB/ABS guidelines for the ethical treatment of animals in behavioral research and teaching and all applicable international, national, and/or institutional guidelines for the care and use of animals were followed. The handling of birds was performed with maximum care and minimal disturbance to nests and the landscape.

