## Supplementary figures for "Golden eagles regularly use gravity waves to soar in the Alps: new insights from high-resolution weather data"

### **Supplementary material**

Tom Carrard[1*], Elham Nourani[2,3,4], Lukas Jansing[5], Tim Zimmermann[1], Petra Sumasgutner[6,7], Matthias Tschumi[8], David Jenny[8], Martin Wikelski[2,3], Kamran Safi[2,3], Michael Sprenger[1†] & Martina Scacco[2,3,4†]

[1]Institute for Atmospheric and Climate Science, ETH Zürich, Zürich, Switzerland

[2]Max Planck Institute of Animal Behavior, Department of Migration, Am Obstberg 1, Radolfzell, 78315, Germany

[3]University of Konstanz, Department of Biology, Universitätsstraße 10, Konstanz, 78464, Germany

[4]Centre for the Advanced Study of Collective Behaviour, University of Konstanz, Universitätsstraße 10, Konstanz, 78464, Germany

[5]Federal Office of Meteorology and Climatology MeteoSwiss, Zurich, Switzerland

[6]Konrad Lorenz Research Center (KLF), Core Facility for Behavior and Cognition, Austria

[7]University of Vienna, Department of Behavioral and Cognitive Biology, Grünau/Almtal, Austria

[8]Swiss Ornithological Institute, Sempach, Switzerland

[†]These authors contributed equally to the study.

### 1 Gravity wave case studies

We present here two additional cases of striking use of a gravity wave for soaring flight.

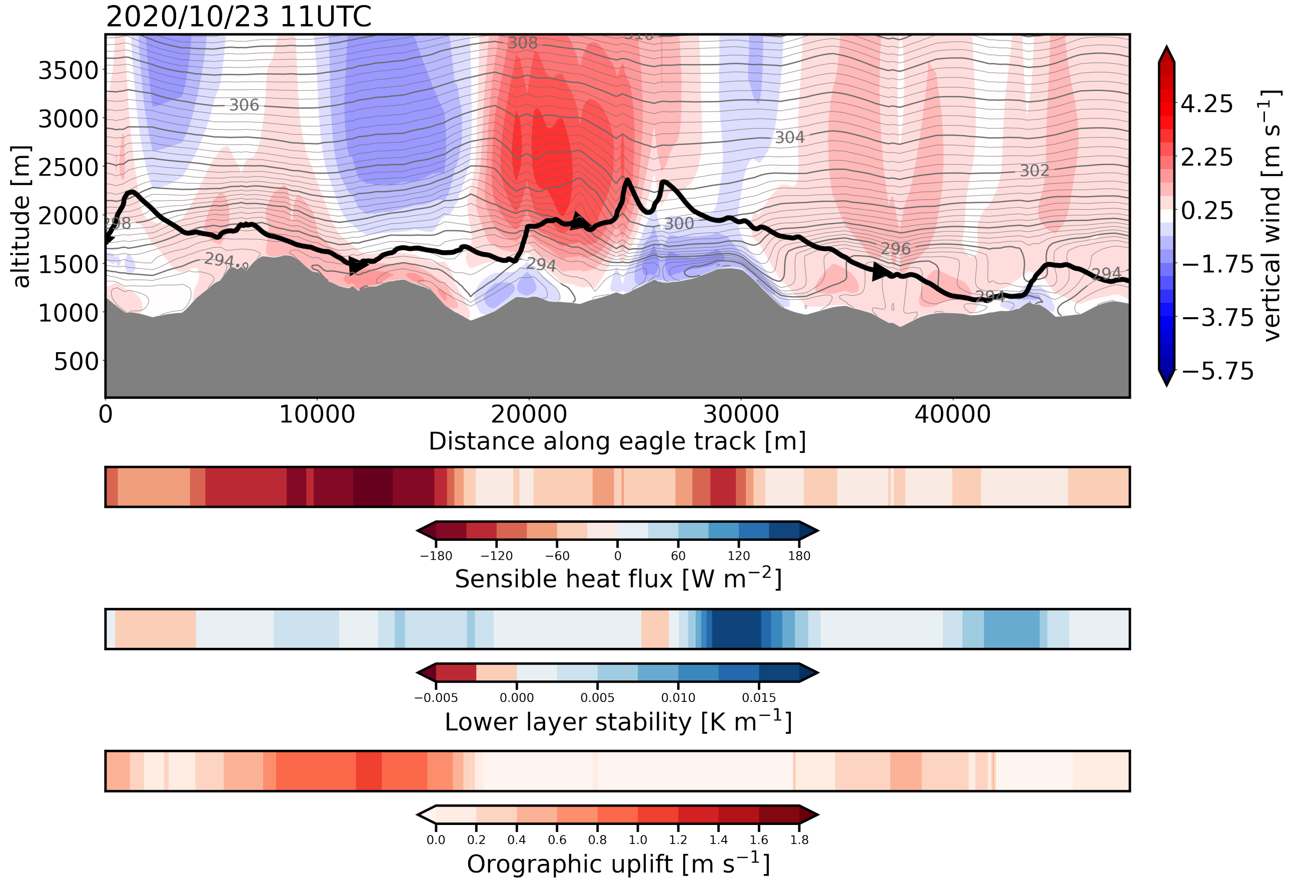


Figure 1: As in fig. 6. The gravity wave updraught corresponds to the large region of vertical wind speed around 20 km. From the the start of the updraught (right before 20 km) to its end, the eagle is flying in a succession of soaring (steep vertical ascent) and gliding (flat or descending segment) and gains overall about 800 metres. While some small-scale thermals may not be excluded near the surface (relatively high sensible heat flux), the eagle is flying far from the ground in a strongly stratified region (lines of potential temperatures are very close together). This suggests that the large ascent is fuelled by the gravity wave updraught.

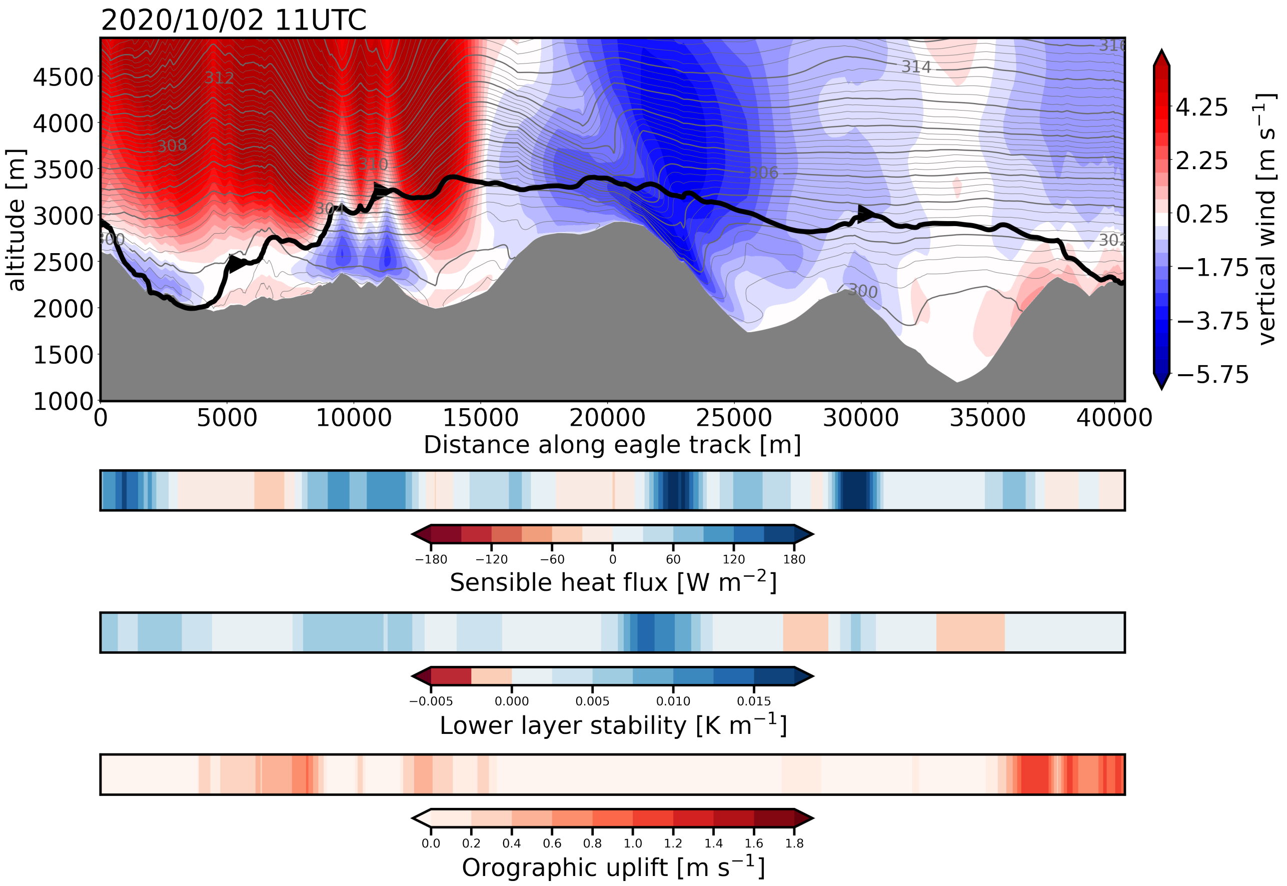


Figure 2: As in fig. 6. The gravity wave updraught corresponds to the large region of high vertical velocity spanning the first 15 km. The vertical velocities are substantial, exceeding $5\,\mathrm{m\,s^{-1}}$. The eagle may first benefit of local orographic lifting (around 5 km) that allows it to reach the larger gravity wave updraught at higher altitude. Between 6 km and 15 km, the eagle is lifted by the gravity wave updraught and gains about 600 metres. It then glides down in the subsequent region of negative vertical velocity.

#### 2 Updraught classification

We provide three additional examples of how the labelling of the updraughts was performed.

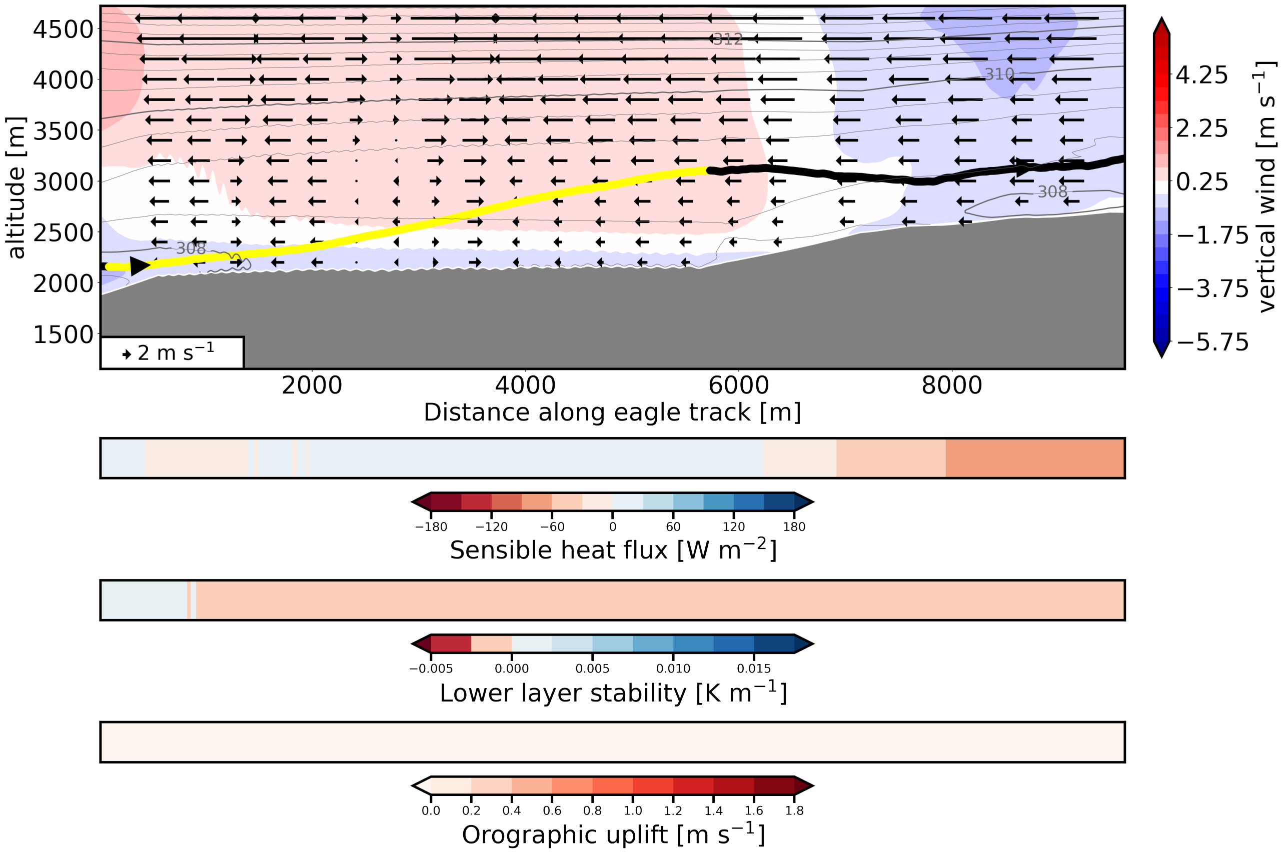


Figure 3: Example of a soaring segment classified as gravity wave updraught. The figure depicts the vertical profiles of vertical wind speed (shading), potential temperature (grey contours) and the horizontal wind projected along the eagle trajectory (arrows). The grey shading represents topography. The lower panels show the sensible heat flux (negative values for an upward flux), lower layer stability (potential temperature gradient in the lower 200 m), and orographic lifting proxy. The black line shows the eagle track and the yellow part corresponds to the labelled soaring segment (larger arrows show the direction of flight). The gravity wave is the large region of positive vertical velocity present over most of the eagle track. The low surface sensible heat flux allows to exclude thermal updraught, while the low orographic lifting proxy indicates that the vertical wind field is not induced by local lifting. The first part of the ascent is probably not powered by the gravity wave, as it is unlikely that the updraught reaches as close to the surface. The eagle may be flapping or benefiting from local turbulence to reach the large gravity wave updraught.

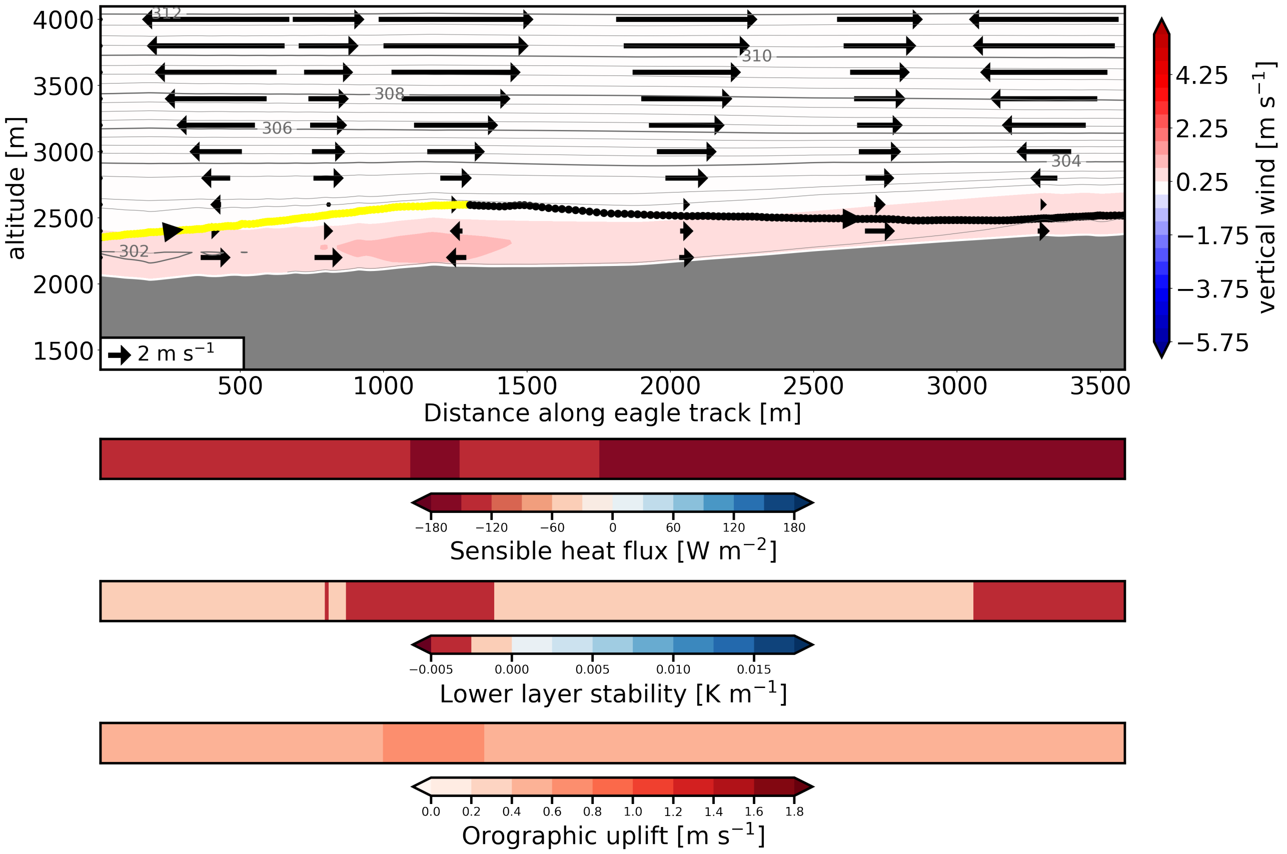


Figure 4: Example of a soaring segment classified as 'thermal/orographic lifting' updraught. The figure depicts the vertical profiles of vertical wind speed (shading), potential temperature (grey contours) and the horizontal wind projected along the eagle trajectory (arrows). The grey shading represents topography. The lower panels show the sensible heat flux (negative values for an upward flux), lower layer stability (potential temperature gradient in the lower 200 m), and orographic lifting proxy. The black line shows the eagle track and the yellow part corresponds to the labelled soaring segment (larger arrows show the direction of flight). From the strong winds at upper level and the relatively high orographic lifting proxy, one can deduce that orographic lifting is occurring and probably contributing to fuel the eagle soaring segment. However, the high sensible heat flux and relatively weak stratification (near the surface) does not allow to exclude thermal formation, such that the soaring segment is probably fuelled by both dynamic and thermal effects.

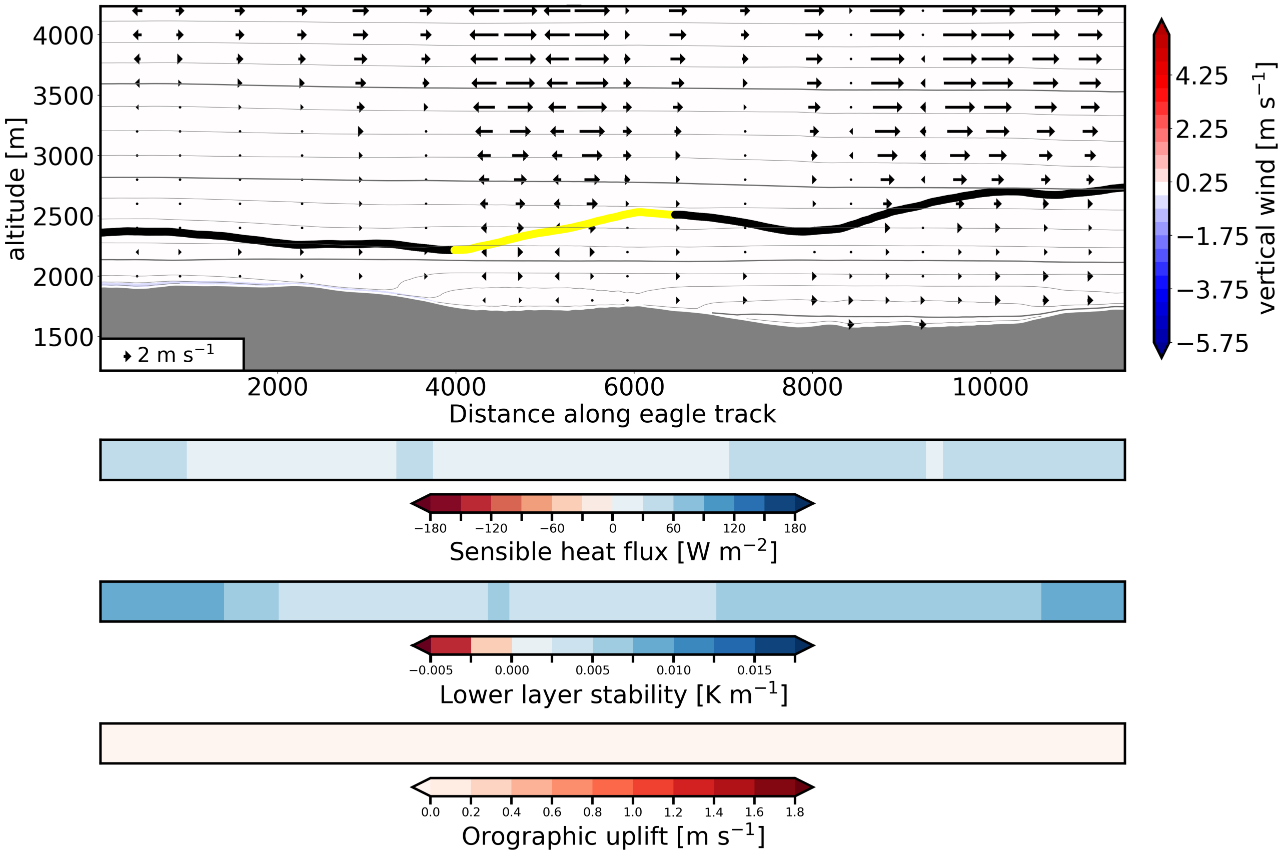


Figure 5: Example of a soaring segment classified as 'unknown'. The figure depicts the vertical profiles of vertical wind speed (shading), potential temperature (grey contours) and the horizontal wind projected along the eagle trajectory (arrows). The grey shading represents topography. The lower panels show the sensible heat flux (negative values for an upward flux), lower layer stability (potential temperature gradient in the lower 200 m), and orographic lifting proxy. The black line shows the eagle track and the yellow part corresponds to the labelled soaring segment (larger arrows show the direction of flight). The soaring segment is located in a region where no wave is modelled, no apparent orographic lifting is present, and the surface sensible heat flux are indicative of a low probability of thermal formation. In this context, we cannot attribute the segment to any of the categories without making strong assumptions. For this reason, the soaring segment is left as 'unknown'.
